# Prospective randomized controlled study directly comparing tadalafil and tamsulosin for male patients with lower urinary tract symptoms

**DOI:** 10.1101/404566

**Authors:** Takumi Takeuchi, Satoshi Toyokawa, Yumiko Okuno, Noriko Ishida, Masanari Yamagoe, Masayoshi Zaitsu, Koji Mikami

**Affiliations:** Department of Urology, Kanto Rosai Hospital, 1-1 Kizukisumiyoshi-cho, Nakahara-ku, Kawasaki 211-8510, Japan; Department of Public Health, Graduate School of Medicine, The University of Tokyo, 7-3-1 Hongo, Bunkyo-ku, Tokyo 113-0033, Tokyo, Japan; Department of Pharmacy, Kanto Rosai Hospital, 1-1 Kizukisumiyoshi-cho, Nakahara-ku, Kawasaki 211-8510, Japan; Yamagoe Urology Clinic, 3-252 Kosugi-cho, Nakahara-ku, Kawasaki 211-0063, Japan

**Keywords:** BPH, LUTS, tadalafil, tamsulosin

## Abstract

Lower urinary tract symptoms are widespread in elderly men and often suggestive of benign prostatic hyperplasia (LUTS/BPH). A randomized, prospective, and open-labeled trial directly comparing the effects of tadalafil (a phosphodiesterase 5 inhibitor) 5 mg once daily and tamsulosin (an α1-blocker) 0.2 mg once daily for 12 weeks in LUTS/BPH patients was conducted. Data were recorded before randomization as well as at 4, 8, and 12 weeks after medication. Fifteen patients allocated tadalafil and 20 allocated tamsulosin completed 12 weeks of medication. Total IPSS, IPSS voiding, and IPSS-QOL scores declined with medication, but there was no difference between drugs. IPSS storage scores reduced more in the tamsulosin group than tadalafil group. OABSS did not decline significantly with medication. IIEF5 was maintained in the tadalafil group, but declined in the tamsulosin group. The maximum flow rate and post-void residual urine volume did not significantly change with medication. Daytime, night-time, and 24-hour urinary frequencies as well as the mean and largest daytime, night-time, and 24-hour voiding volumes per void did not significantly change with medication. In conclusion, tamsulosin was more effective to reduce storage symptoms than tadalafil. Tadalafil had the advantage of maintaining the erectile function.

## Introduction

Lower urinary tract symptoms (LUTS) are common in elderly men and often suggestive of benign prostatic hyperplasia (LUTS/BPH). Pharmacological management of LUTS/BPH normally includes α1-blockers, 5α-reductase inhibitors, and phosphodiesterase 5 inhibitors. α1-Blockers and phosphodiesterase 5 inhibitors reduce smooth muscle tone in the prostate and urethra by inhibiting the effect of catecholamines and increasing intracellular cyclic guanosine monophosphate, respectively. Thus, they alleviate bladder outlet obstruction. 5α-Reductase inhibitors inhibit 5α-reductase, a nuclear-bound steroid enzyme that converts testosterone into a more potent form, dihydrotestosterone (DHT). Reduced DHT in prostatic tissue causes apoptosis of prostate epithelial cells (1) and reduces the prostate volume by as much as 30%.

The administration of tadalafil, a phosphodiesterase 5 inhibitor, to LUTS/BPH patients improved the International Prostate Symptom Score (IPSS) total score (IPSS-T) compared with placebo (2-10) as well as IPSS voiding (IPSS-V) and IPSS storage (IPSS-S) subscores (3, 6, 7, 11), but those positive effects on urination were less marked for elderly patients more than 75 years old (12). In addition, tadalafil improved the IPSS quality of life (QOL) score (2, 6-8, 12). Nevertheless, tadalafil did not enhance the maximum flow rate (MFR) nor reduce the post-void residual urine on uroflowmetry in many studies (2-5, 10), although they were occasionally ameliorated (13, 14).

Combinations of tadalafil and another type of drug for BPH were also investigated. A meta-analysis showed that combined therapy of tadalafil with α1-blockers more markedly improved IPSS, the International Index of Erectile Function (IIEF), and MFR compared with α-blockers alone (13). IPSS-T, IPSS-S, IPSS-V, and IPSS-QOL were improved by a combination of tadalafil and finasteride, a 5α-reductase inhibitor, compared with finasteride alone (15).

Another effect of tadalafil is to improve erectile dysfunction (ED) as evaluated by IIEF (2-5, 10, 16), observed regardless of the severity of concurrent LUTS/BPH (7). On the contrary, the severity of ED reportedly did not influence the improvement of LUTS/BPH (17, 18).

Here, we conducted a randomized, prospective, and open-labeled trial directly comparing the effects of tadalafil with those of tamsulosin, an α1-blocker, on urination in LUTS/BPH patients.

## Patients and Methods

Participants were LUTS/BPH patients visiting Kanto Rosai Hospital and Yamagoe Urology Clinic. At the initial visit, all patients underwent a detailed clinical evaluation including a complete medical history, physical examination, urinalysis, and trans-abdominal ultrasonography. Assessment of IPSS, IPSS-QOL Score, Overactive Bladder Symptom Score (OABSS), IIEF5, uroflowmetry, and post-void residual urine volume measurement was performed before randomization.

In addition, patients were asked to create a 24-hour flow-volume chart (24-h-FVC), stating the time and volume of each void and their bedtime and waking time before randomization. The nocturnal urine volume was defined as the volume of voids between bedtime and waking time plus the first-morning void, while the first-morning void was regarded as a normal diurnal voiding episode. Daytime, night-time, and 24-hour (24-h) urinary frequencies as well as daytime, night, and 24-h mean volumes per void were calculated using 24-h-FVC. The largest daytime and night-time voided volumes were also evaluated.

The exclusion criteria were as follows: (a) taking anti-cholinergic drugs, beta3-stimulating drugs, anti-depressants, and flavoxate hydrochloride; (b) previously taking a 5α-reductase inhibitor and chlormadinone acetate; (c) taking nitric acid drugs and nitric oxide-releasing drugs, and soluble guanylate cyclase stimulating drugs; (d) allergic to tadalafil and tamsulosin; (e) unstable angina, cardiac failure more than NYHA3, uncontrollable arrhythmia, hypotension (systolic blood pressure <90 mmHg or diastolic blood pressure <50 mmHg), and hypertension (resting systolic blood pressure >170 mmHg or resting diastolic blood pressure >100 mmHg); (f) cardiac infarction within 3 months, cerebrovascular disease within 6 months; severe renal failure and hepatic failure; (g) unwilling to participate in this study.

Randomization to treatment groups, tadalafil 5 mg once daily or tamsulosin 0.2 mg once daily for 12 weeks, was conducted at a 1:1 ratio using stratification by the prostate volume (≤50 cc or >50 cc) and age (10-year increments). The allocation was achieved using random permuted blocks of size four. Thereafter, IPSS, IPSS-QOL, OABSS, IIEF5, uroflowmetry, and 24-h-FVC were recorded at 4, 8, and 12 weeks after medication. MFR was determined from uroflowmetry, and the post-void residual urine was measured by BVI6100 (Sysmex, Kobe, Japan). This clinical trial was registered with the UMIN Clinical Trials Registry (UMIN000020362).

## Statistics

The age, prostate volume, and prostate-specific antigen (PSA) in both groups were analyzed by the unpaired t-test. IPSS, QOL, OABSS, IIEF5, MFR, post-void residual urine, daytime, night, and 24-h urinary frequencies, largest daytime and night voiding volumes, and mean daytime, night, and 24-h voiding volumes per void were analyzed by two-way repeated-measure analysis of variance (two-way repeated-measure ANOVA) with the drug a fixed effect and duration of either drug administration a random effect. Dunnett’s test and the Welch t-test were conducted as post-hoc tests of the medication duration and drug, respectively. Adverse events were asessed by the two-tailed Fisher’s exact test. Statistical analysis was conducted with R version 3.2.4.

## Ethics

The Ethics Committee of Kanto Rosai Hospital approved the study (2015-21). Written informed consent was received from all participants prior to inclusion in the study. We declare that there is no conflict of interest regarding the publication of this paper.

## Results

In total, 39 patients were recruited between January 2016 and April 2018. α1-Blockers for BPH treatment were washed out at least two weeks before randomization in four patients (10.3%; naftopidil, tadalafil, silodosin, and urapidil in one case, respectively). Five patients (12.8%) had previously undergone prostate biopsy without cancer detection. Pelvic MRI was indicated for most patients with elevated serum PSA (>4.0 ng/mL). With randomization, 18 were allocated to tadalafil 5 mg once daily, and 21 patients were allocated to tamsulosin 0.2 mg once daily. Just after randomization, one patient allocated tadalafil withdrew from this study voluntarily and another allocated tamsulosin was excluded due to being diagnosed with prostate cancer. One patient stopped taking tadalafil after two weeks because of a headache and another taking tadalafil developed urinary retention, necessitating bladder catheter indwelling after four weeks, and so they were excluded from the study. One patient taking tadalafil felt palpitations, but continued medication. Therefore, rates of adverse events were 17.6% in the tadalafil group, but 0% in the tamsulosin group (p=0.09). Thus, 15 allocated tadalafil (88.2%) and 20 patients allocated tamsulosin (95.2%) completed 12 weeks of medication.

Patient characteristics are presented in Table 1. The age, prostate volume, and PSA were not significantly different between tadalafil and tamsulosin groups including all who commenced medication, although the prostate volume in the former was non-significantly smaller than that in the latter.

**Table 1:** Patient characteristics

|  | Tadalafil (n=17) | Tamsulosin (n=20) | p-value |
| --- | --- | --- | --- |
| Age (years) | 67.5±8.5 | 67.1±9.9 | 0.905 |
| Prostate volume (mL) | 38.6±26.1 | 48.1±18.0 | 0.196 |
| PSA (ng/mL) | 4.3±4.3 | 4.3±3.3 | 0.960 |

values: mean ± standard deviation,

### Questionnaires

IPSS-T significantly declined, but there was no difference between drugs (Table 2 and Figure 1). IPSS-V marginally diminished, but again there was no difference between drugs (Table 2 and Figure 2). IPSS-S diminished more in the tamsulosin group than in the tadalafil group (Table 2 and Figure 3). The IPSS-QOL score significantly declined with medication, but there was no difference between drugs (Table 2 and Figure 4). The IPSS-Q7 (nocturia) score did not significantly change with medication (Table 2). OABSS did not decline significantly with medication, and there was no difference between drugs (Table 2 and Figure 5). Baseline (before medication) scores of IIEF5 were unbalanced between tadalafil and tamsulosin groups (6.0±1.5 vs. 8.8±1.4, respectively; mean ± standard error); thus, changes from baseline scores were evaluated. IIEF5 was maintained in the tadalafil group, but declined in the tamsulosin group. ANOVA showed a significant difference between the two drug groups (Table 2 and Figure 6).

**Table 2:**
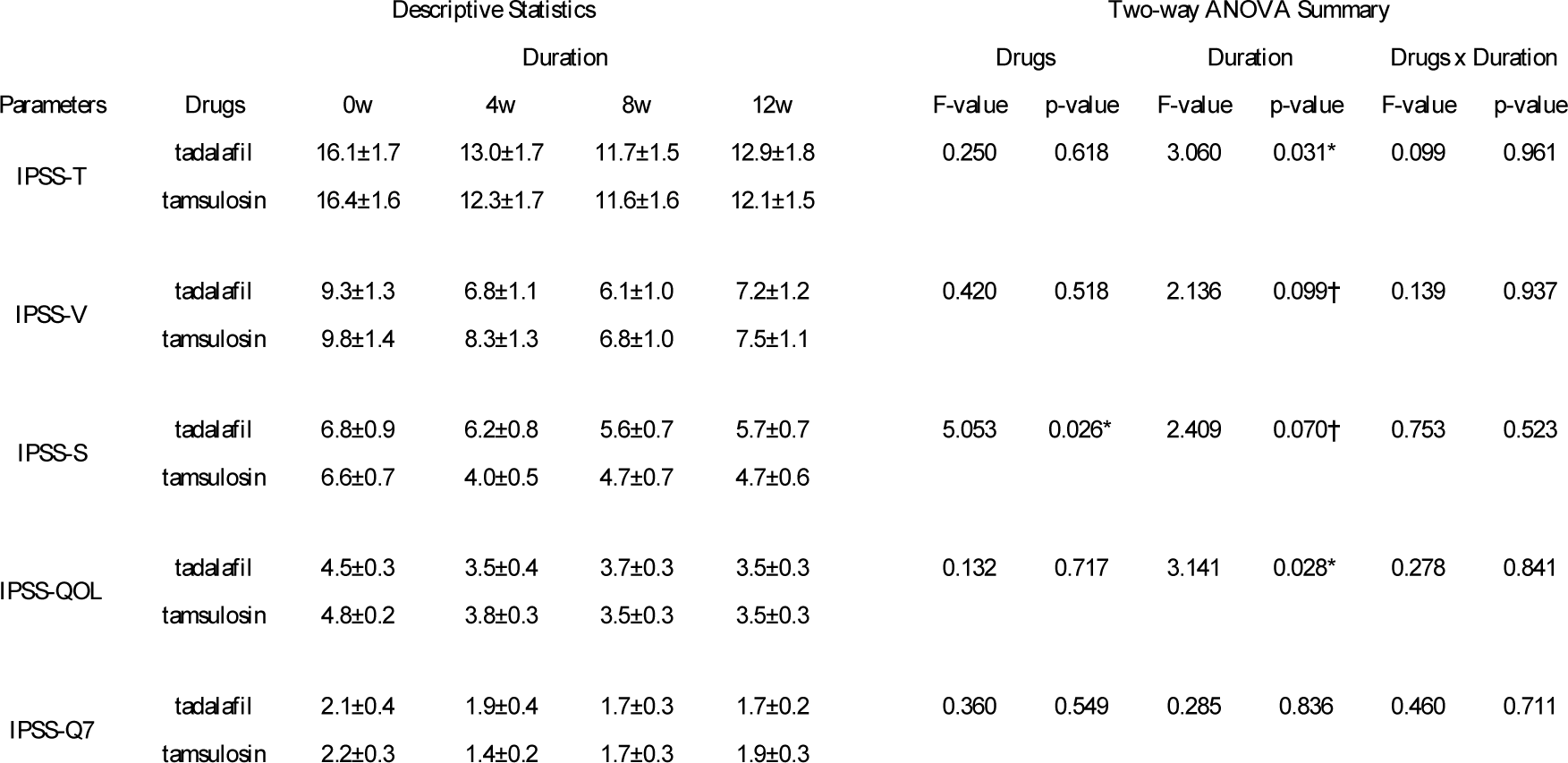
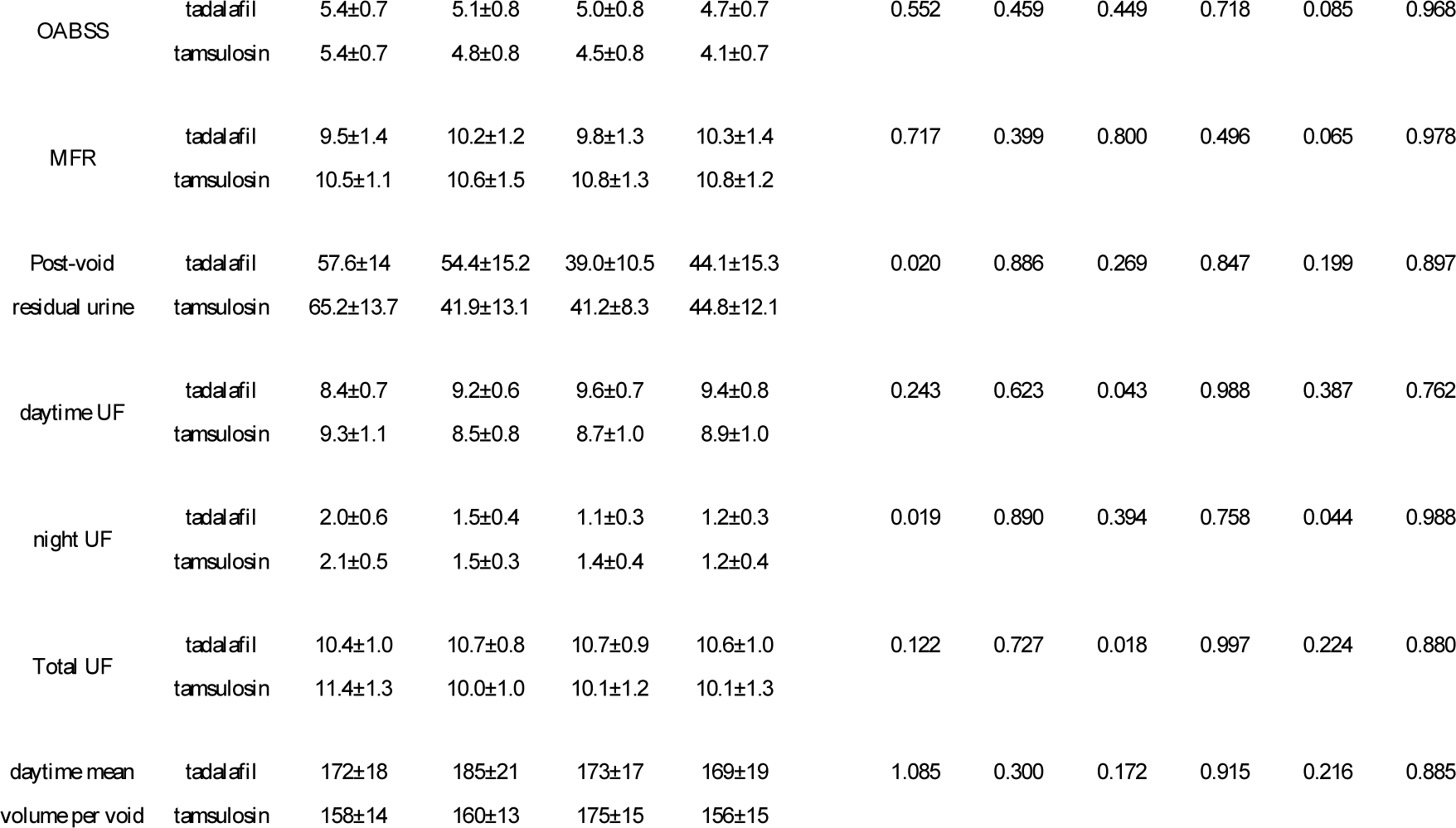
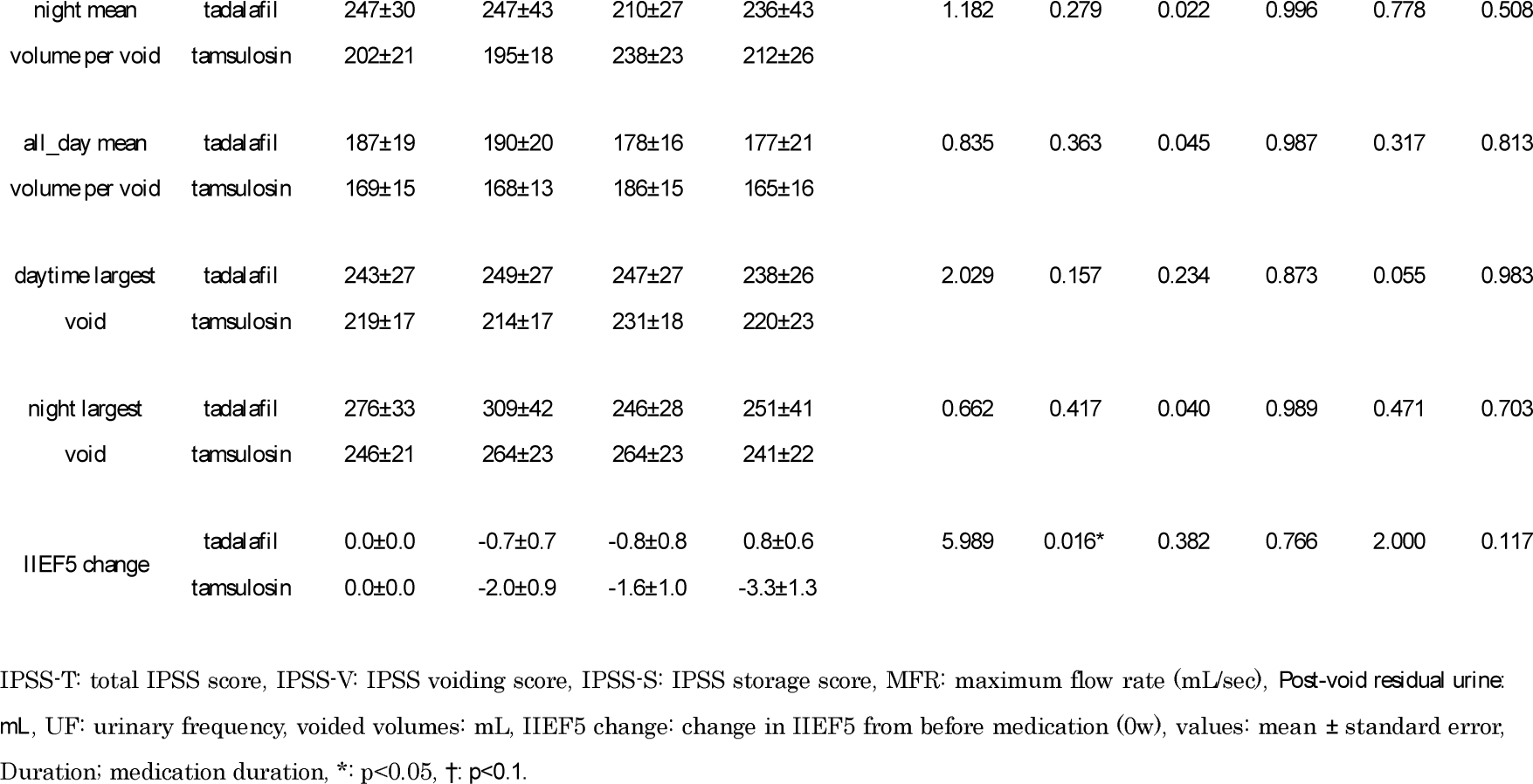
Comparison between tadalafil and tamsulosin administrations.

**Figure 1:**
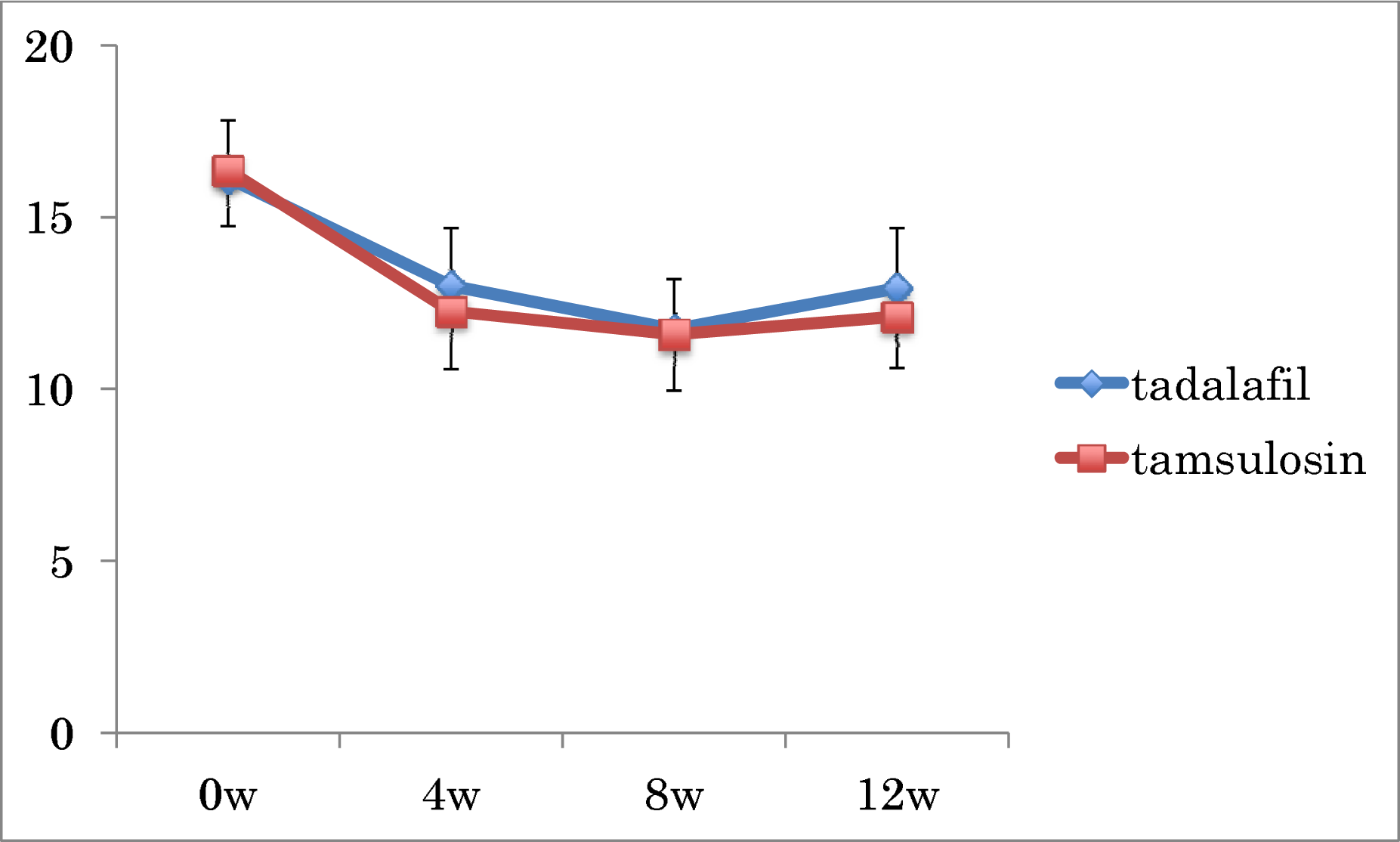
Changes in total IPSS scores. ANOVA showed a significant difference in the medication duration, while there was no difference between drugs.

**Figure 2:**
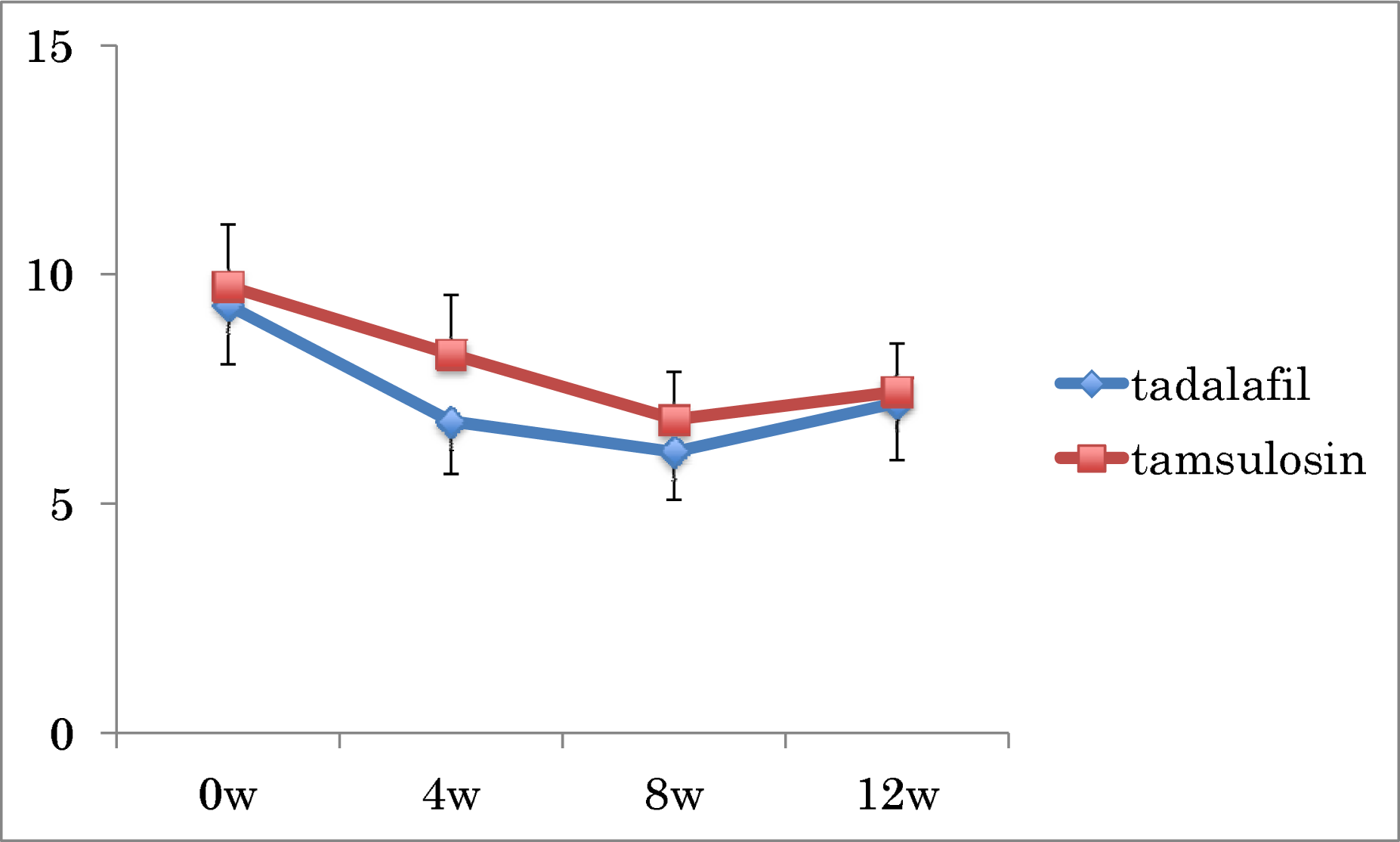
Changes in IPSS voiding scores. ANOVA showed a marginal difference in the medication duration, while there was no difference between drugs.

**Figure 3:**
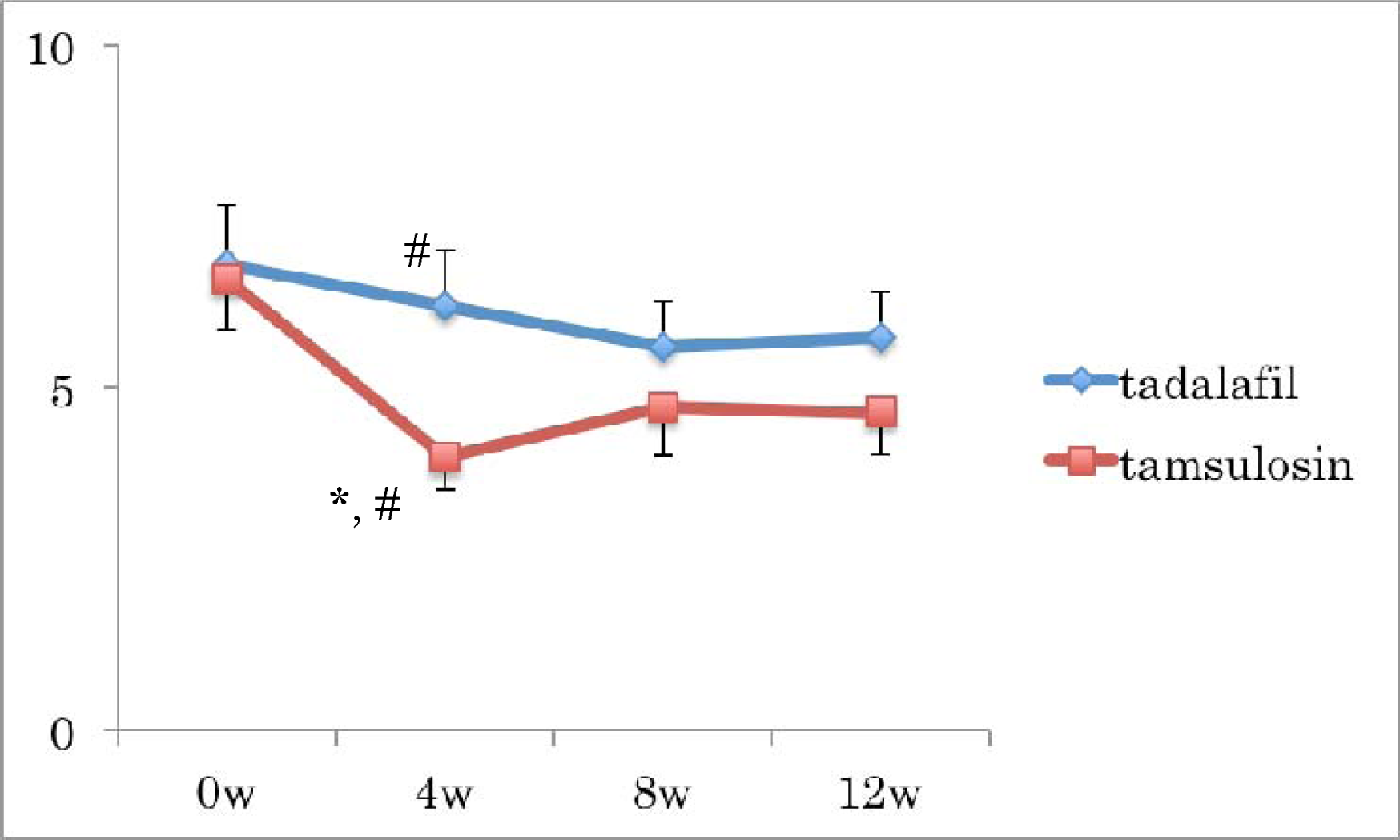
Changes in IPSS storage scores. ANOVA showed a marginal difference in the medication duration and significant difference between drugs. #: p<0.05 between drugs by Welch t-test.

**Figure 4:**
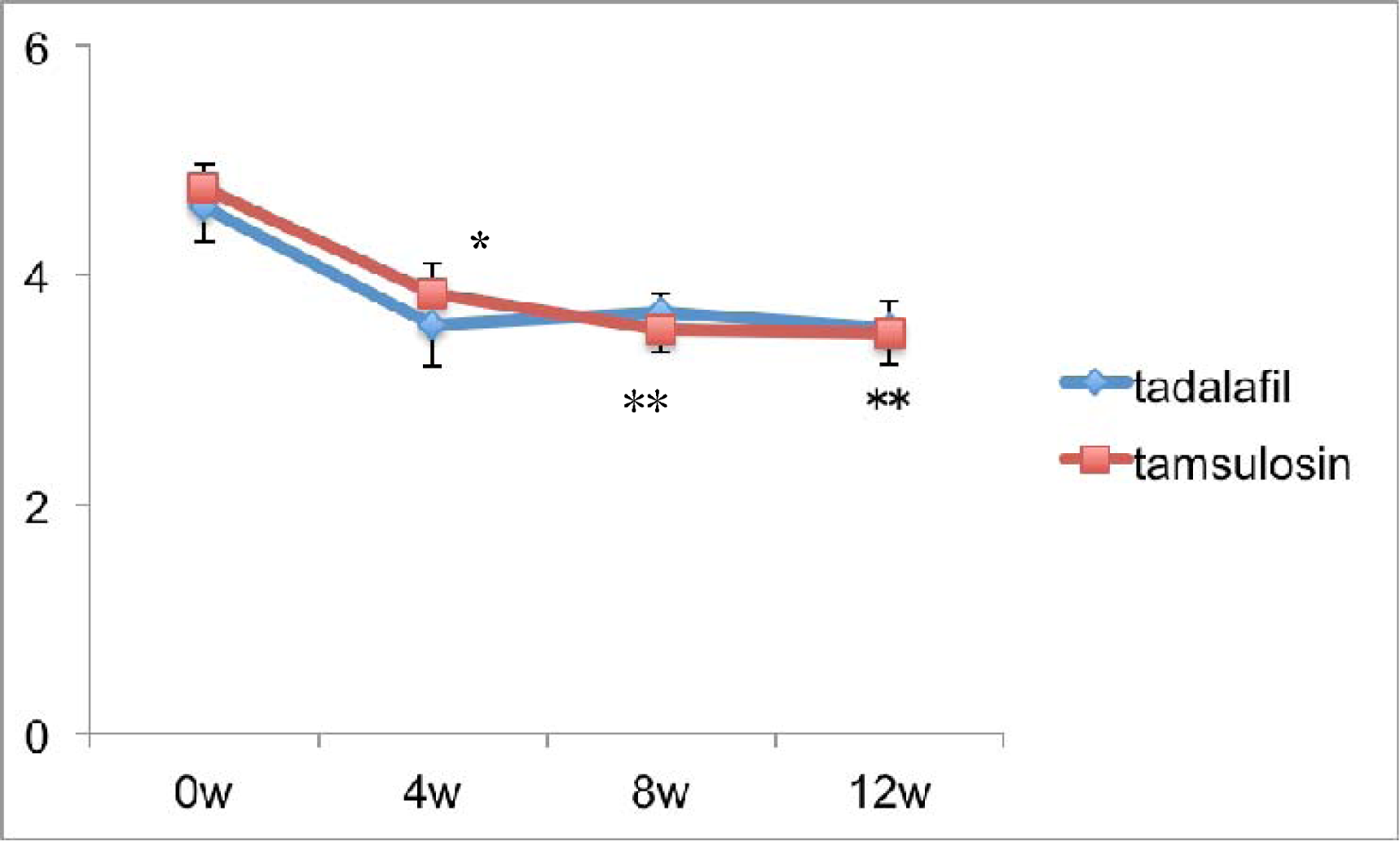
Changes in IPSS QOL scores. ANOVA showed a significant difference in the medication duration, while there was no difference between drugs. *: p<0.05, **: p<0.01 compared with before medication (0w) by Dunnett’s test.

**Figure 5:**
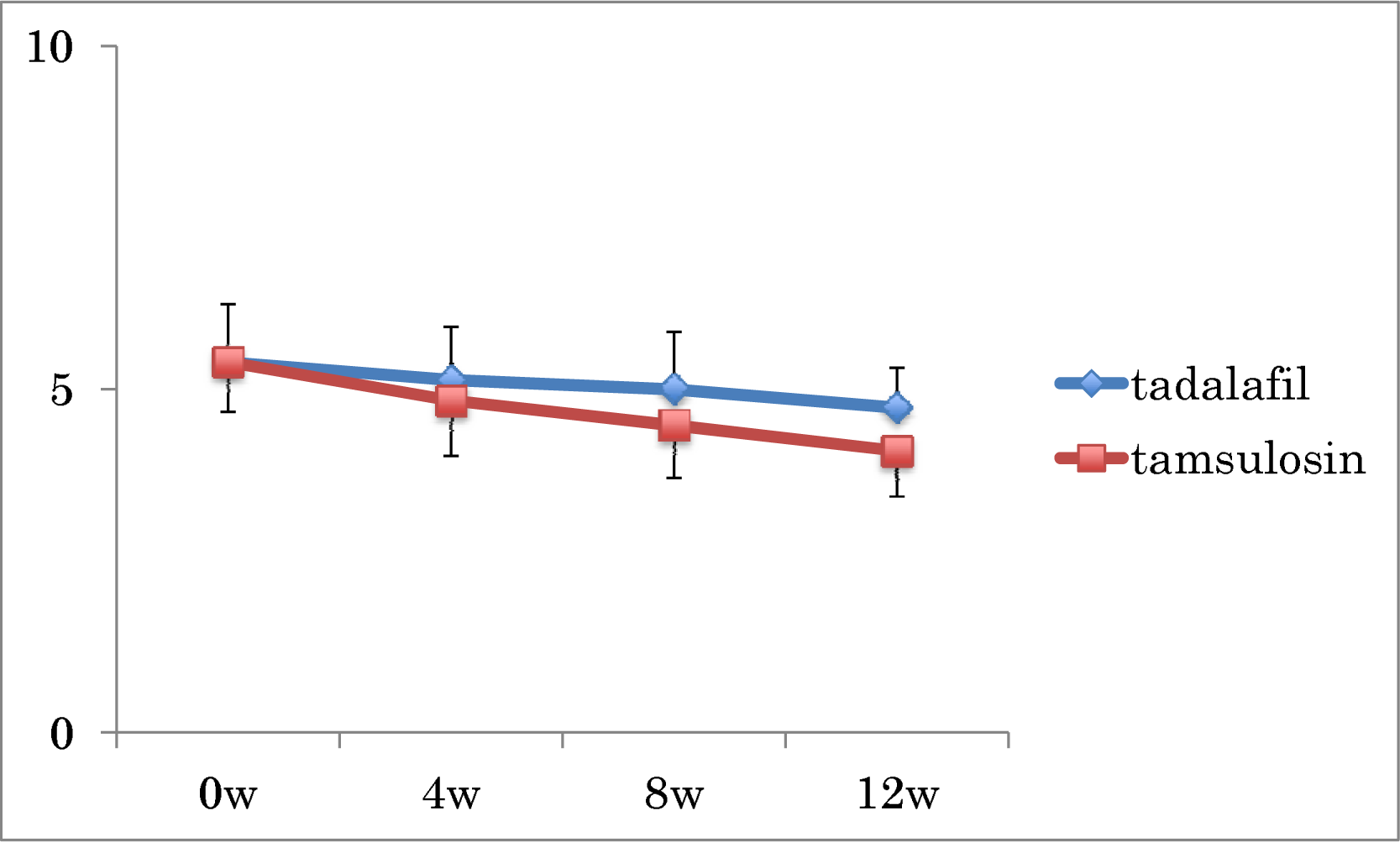
Changes in OABSS scores. ANOVA showed no difference in the medication duration or between drugs.

**Figure 6:**
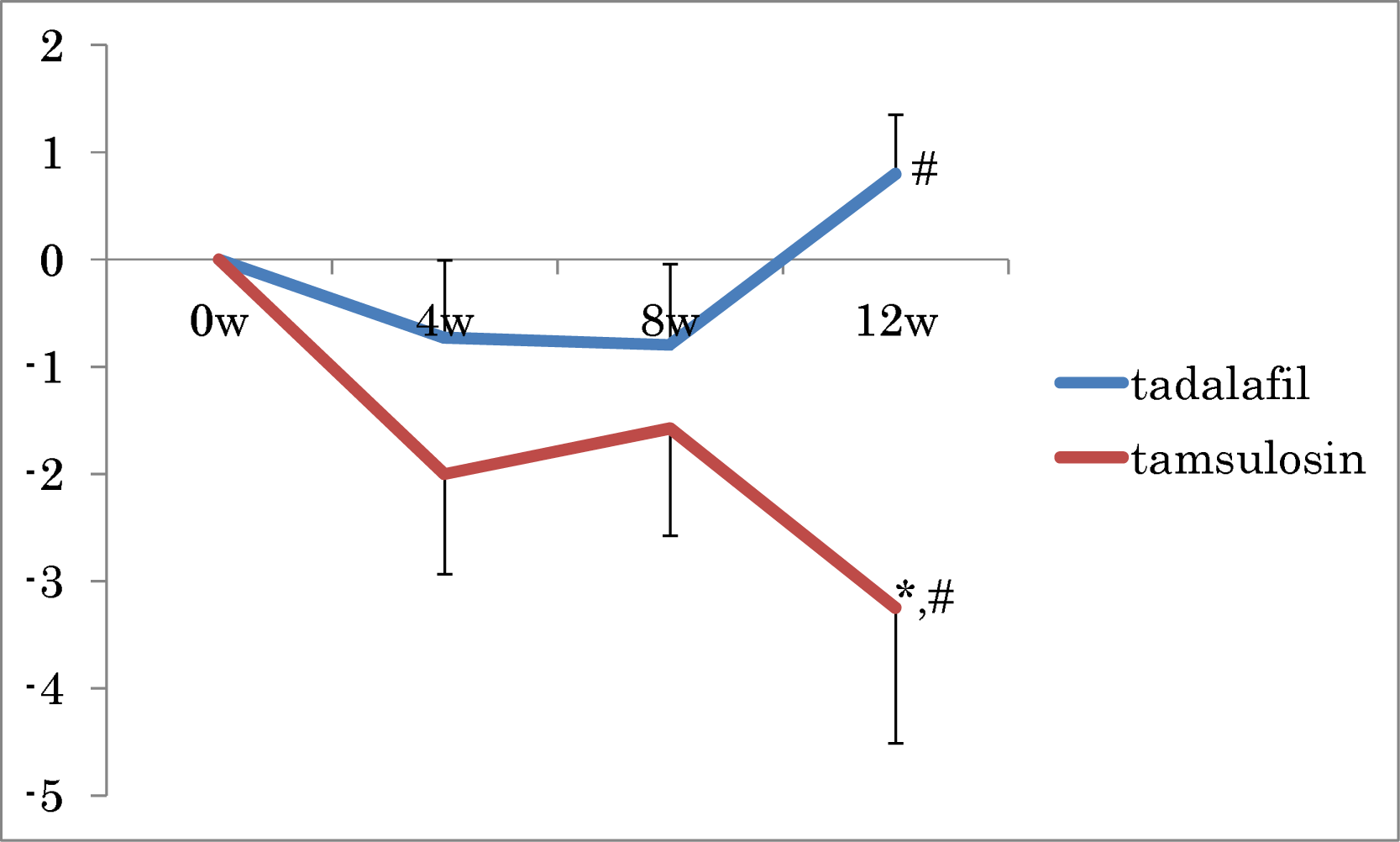
Changes in IIEF5 scores from before medication. *: p<0.05 compared with before medication (0w) by Dunnett’s test. #: p<0.05 between drugs by Welch t-test.

### Uroflowmetry parameters

The maximum flow rate and post-void residual urine volume did not significantly change with medication (Table 2).

### 24-h-FVC parameters

As shown in Table 2, daytime, night-time, and 24-h urinary frequencies did not significantly change with medication, and there were no differences between drugs. In addition, the largest daytime and night voiding volumes as well as mean daytime, night-time, and 24-h voiding volumes per void did not significantly change with medication, and there were no differences between drugs.

For all parameters investigated, ANOVA demonstrated the lack of a significant interaction between the drugs and medication duration.

## Discussion

Comparison of α-blockers and tadalafil is significant in order to choose drugs for the medical treatment of BPH. In 2012, Yokoyama et al. prospectively randomized 612 LUTS/BPH Asian patients into four groups, each of which was administered placebo, 0.2 mg of tamsulosin once daily, 2.5 mg of tadalafil once daily, or 5 mg of tadalafil once daily for 12 weeks (6). They concluded that both tadalafil and tamsulosin reduced IPSS compared with the placebo. However, they did not directly and statistically compare the two drugs for unknown reasons. Judging from the data they showed, reductions in total IPSS, IPSS-V, and IPSS-QOL scores were greater in the tamsulosin group than tadalafil group, while IPSS-S was comparable between the groups.

Similarly, Oelke et al. prospectively randomized 511 LUTS/BPH patients mainly in European countries into three groups, administered a placebo, 0.4 mg of tamsulosin once daily, or 5 mg of tadalafil once daily for 12 weeks (13). They also concluded that both tadalafil and tamsulosin reduced IPSS compared with the placebo. They also failed to conduct a direct and statistical comparison between tadalafil and tamsulosin groups. Judging from the data they showed, reductions in IPSS-T, IPSS-V, and IPSS-QOL scores were greater in the tadalafil group than tamsulosin group, being different from the Asian population, while IPSS-S was comparable between the groups.

The present study enrolled a small number of LUTS/BPH patients. Nevertheless, ANOVA revealed that IPSS-T, IPSS-S, IPSS-V, and IPSS-QOL were reduced with active treatments by tadalafil and tamsulosin. Additionally, tamsulosin significantly reduced IPSS-S more than tadalafil. It is difficult to know whether this result is consistent with the studies done by Yokoyama or Oelke, because they did not directly compare between the two drugs.

Significant changes in uroflowmetry and 24-h-FVC parameters, OABSS, and IPSS-Q7 were not detected by ANOVA, thus neither tadalafil nor tamsulosin had a significant influence on those factors. The finding that tadalafil does not improve uroflowmetry parameters is in line with most previous reports, while the finding that tadalafil does not have significant effects on 24-h-FVC parameters is relatively novel.

Yoshida et al. prospectively randomized 188 LUTS/BPH patients into two groups: 5 mg of tadalafil once daily and 4 mg of silodosin (an α1-blocker) twice daily for 8 weeks. They showed that reductions in IPSS-T, IPSS-S, IPSS-QOL, and OABSS scores were more marked in the silodosin group than in the tadalafil group, while IPSS-V was comparable. In addition, MFR tended to increase more with silodosin than tadalafil in their study. Their results were essentially in line with those of this study. α1-Blockers may alleviate storage symptoms of LUTS/BPH patients more than tadalafil (19).

As previously reported, tadalafil had more favorable effects on ED compared with tamsulosin. In this study, adverse events were slightly more frequent in the tadalafil group than tamsulosin group, without significance. Drugs were well-tolerated in both groups. Limitations of this study include the small number of patients from a few institutions and that it was an open-labeled trial. Therefore, weaker effects of either drug might have gone undetected. In conclusion, tamsulosin was more effective to reduce storage symptoms than tadalafil in LUTS/BPH patients, while voiding symptoms, IPSS-T, IPSS-QOL were comparably declined with the drugs. Tadalafil had the advantage maintaining the erectile function.

## References

1. Andriole G, Bruchovsky N, Chung LW, Matsumoto AM, Rittmaster R, Roehrborn C, Russell D, Tindall D. Dihydrotestosterone and the prostate: the scientific rationale for 5alpha-reductase inhibitors in the treatment of benign prostatic hyperplasia. J Urol. 172: 1399–403, 2004.

2. McVary KT, Roehrborn CG, Kaminetsky JC, Auerbach SM, Wachs B, Young JM, Esler A, Sides GD, Denes BS. Tadalafil relieves lower urinary tract symptoms secondary to benign prostatic hyperplasia. J Urol. 177: 1401–7, 2007.

3. Roehrborn CG, McVary KT, Elion-Mboussa A, Viktrup L. Tadalafil administered once daily for lower urinary tract symptoms secondary to benign prostatic hyperplasia: a dose finding study. J Urol. 180: 1228–34, 2008.

4. Porst H, Kim ED, Casabe AR, Mirone V, Secrest RJ, Xu L, Sundin DP, Viktrup L; LVHJ study team. Efficacy and safety of tadalafil once daily in the treatment of men with lower urinary tract symptoms suggestive of benign prostatic hyperplasia: results of an international randomized, double-blind, placebo-controlled trial. Eur Urol. 60: 1105–13, 2011.

5. Gacci M, Corona G, Salvi M, Vignozzi L, McVary KT, Kaplan SA, Roehrborn CG, Serni S, Mirone V, Carini M, Maggi M. A systematic review and meta-analysis on the use of phosphodiesterase 5 inhibitors alone or in combination with a -blockers for lower urinary tract symptoms due to benign prostatic hyperplasia. Eur Urol. 61: 994–1003, 2012.

6. Yokoyama O, Yoshida M, Kim SC, Wang CJ, Imaoka T, Morisaki Y, Viktrup L. Tadalafil once daily for lower urinary tract symptoms suggestive of benign prostatic hyperplasia: a randomized placebo- and tamsulosin-controlled 12-week study in Asian men. Int J Urol. 20: 193–201, 2013.

7. Porst H, Roehrborn CG, Secrest RJ, Esler A, Viktrup L. Effects of tadalafil on lower urinary tract symptoms secondary to benign prostatic hyperplasia and on erectile dysfunction in sexually active men with both conditions: analyses of pooled data from four randomized, placebo-controlled tadalafil clinical studies. J Sex Med. 10: 2044–52, 2013.

8. Porst H, Oelke M, Goldfischer ER, Cox D, Watts S, Dey D, Viktrup L. Efficacy and safety of tadalafil 5 mg once daily for lower urinary tract symptoms suggestive of benign prostatic hyperplasia: subgroup analyses of pooled data from 4 multinational, randomized, placebo-controlled clinical studies. Urology. 82: 667–73, 2013.

9. Nickel JC, Brock GB, Herschorn S, Dickson R, Henneges C, Viktrup L. roportion of tadalafil-treated patients with clinically meaningful improvement in lower urinary tract symptoms associated with benign prostatic hyperplasia - integrated data from 1,499 study participants. BJU Int. 115: 815–21, 2015.

10. Gacci M, Andersson KE, Chapple C, Maggi M, Mirone V, Oelke M, Porst H, Roehrborn C, Stief C, Giuliano F. Latest Evidence on the Use of Phosphodiesterase Type 5 Inhibitors for the Treatment of Lower Urinary Tract Symptoms Secondary to Benign Prostatic Hyperplasia. Eur Urol. 70: 124–133, 2016.

11. Chapple CR, Roehrborn CG, McVary K, Ilo D, Henneges C, Viktrup L. Effect of tadalafil on male lower urinary tract symptoms: an integrated analysis of storage and voiding international prostate symptom subscores from four randomised controlled trials. Eur Urol. 67: 114–122, 2015.

12. Oelke M, Wagg A, Takita Y, Buttner H, Viktrup L. Efficacy and safety of tadalafil 5 mg once daily in the treatment of lower urinary tract symptoms associated with benign prostatic hyperplasia in men aged ≥75 years: integrated analyses of pooled data from multinational, randomized, placebo-controlled clinical studies. BJU Int. 119: 793–803, 2017.

13. Oelke M, Giuliano F, Mirone V, Xu L, Cox D, Viktrup L. Monotherapy with tadalafil or tamsulosin similarly improved lower urinary tract symptoms suggestive of benign prostatic hyperplasia in an international, randomised, parallel, placebo-controlled clinical trial. Eur Urol. 61: 917–25, 2012.

14. Roehrborn CG, Chapple C, Oelke M, Cox D, Esler A, Viktrup L. Effects of tadalafil once daily on maximum urinary flow rate in men with lower urinary tract symptoms suggestive of benign prostatic hyperplasia. J Urol. 191: 1045–50, 2014.

15. Casabe A, Roehrborn CG, Da Pozzo LF, Zepeda S, Henderson RJ, Sorsaburu S, Henneges C, Wong DG, Viktrup L. Efficacy and safety of the coadministration of tadalafil once daily with finasteride for 6 months in men with lower urinary tract symptoms and prostatic enlargement secondary to benign prostatic hyperplasia. J Urol. 191: 727–33, 2014.

16. Giuliano F, Oelke M, Jungwirth A, Hatzimouratidis K, Watts S, Cox D, Viktrup L. Tadalafil once daily improves ejaculatory function, erectile function, and sexual satisfaction in men with lower urinary tract symptoms suggestive of benign prostatic hyperplasia and erectile dysfunction: results from a randomized, placebo- and tamsulosin-controlled, 12-week double-blind study. J Sex Med. 10: 857–65, 2013.

17. Brock G, Broderick G, Roehrborn CG, Xu L, Wong D, Viktrup L. Tadalafil once daily in the treatment of lower urinary tract symptoms (LUTS) suggestive of benign prostatic hyperplasia (BPH) in men without erectile dysfunction. BJU Int. 112: 990–7, 2013.

18. Brock GB, McVary KT, Roehrborn CG, Watts S, Ni X, Viktrup L, Wong DG, Donatucci C. Direct effects of tadalafil on lower urinary tract symptoms versus indirect effects mediated through erectile dysfunction symptom improvement: integrated data analyses from 4 placebo controlled clinical studies. J Urol. 191: 405–11, 2014.

19. Yoshida M, Origasa H, Seki N. Comparison of silodosin versus tadalafil in patients with lower urinary tract symptoms associated with benign prostatic hyperplasia. Low Urin Tract Symptoms. 9: 176–186, 2017.

